# Predicting Phage–Host Interactions Across Taxonomic Levels: A Systematic Review and Meta-Analysis for Microbial Ecology

**DOI:** 10.64898/2026.04.28.721508

**Authors:** Danitza Xiomara Romero-Calle, Miguel Yucra Rojas, Mathias Middelboe

**Affiliations:** Instituto de Investigaciones Industriales, Ingeniería Industrial, Facultad de Ingeniería, Universidad Mayor de San Andrés, La Paz, Bolivia; Marine Biological Section, Department of Biology, University of Copenhagen, Copenhagen, Denmark; HADAL & Nordcee, Department of Biology, University of Southern Denmark, Odense, Denmark

**Keywords:** Bacteriophage, bacteria, phage-host interaction, machine learning

## Abstract

The prediction of phage-host interactions is key for several applications in biotechnology, medicine, and microbial ecology. Wide studies in machine learning tools have allowed the exploration of these interactions across multiple taxonomic levels. A systematic review and meta-analysis were conducted on 570 records retrieved from PubMed, Scopus, and Web of Science. Eleven studies were selected for the meta-analysis, encompassing 61 datasets. Precision across taxonomic levels (Domain, Phylum, Class, Order, Family, Genus, Species) was evaluated for several prediction tools. Statistical tests, including the Shapiro-Wilk and ANOVA tests, were used. A mixed-effects meta-regression model was used to examine the impact of taxonomic subgroups on the prediction of the proportion of Correctly Predicted PHIs. The results indicated significant variability in the performance of prediction tools across taxonomic levels. Domain-level predictions exhibited near-perfect Proportion of Correctly Predicted PHIs (0.99), whereas finer resolutions (Family and Order) showed considerable variability, with average precision values of 0.682 and 0.775, respectively. The mixed-effects meta-regression analysis revealed that Family and Species taxonomic subgroups were associated with significant reductions in the prediction Proportion of Correctly Predicted PHIs with effect sizes of −0.1464 and −0.1944, respectively. Residual heterogeneity was negligible, indicating that the moderators adequately explained the variability in prediction precision. This study highlights the importance of selecting the appropriate prediction tool based on the desired taxonomic resolution. The findings emphasize the need for further refinement of prediction algorithms, particularly at the Family and Species levels, where tools exhibit the most variability.

Graphical Abstract.
Overview of the systematic review and meta-analysis framework evaluating ML-based phage–host interaction prediction tools across taxonomic levels.

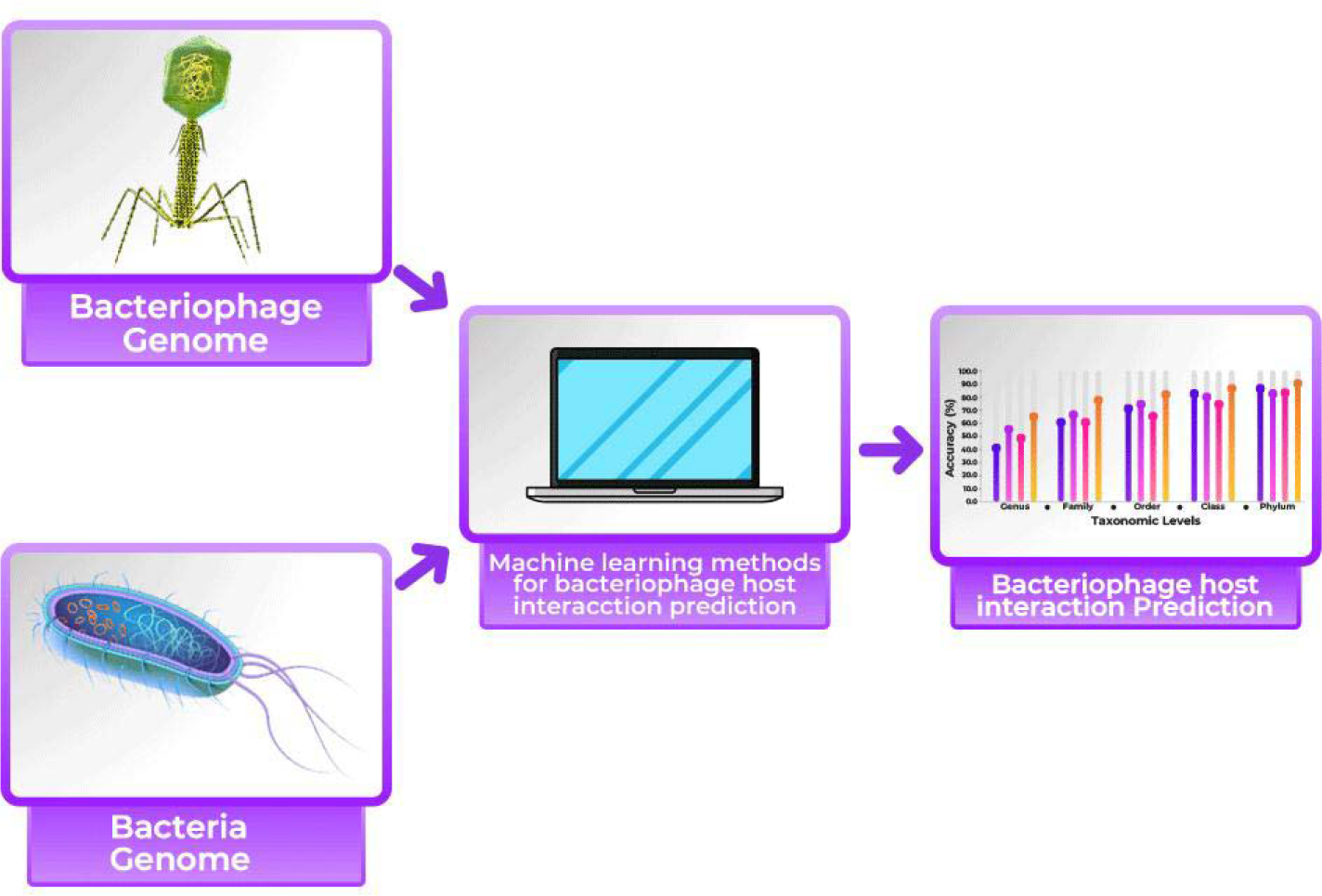

## Introduction

Bacteriophages are emerging as promising alternatives to antibiotics due to increasing antimicrobial resistance. Due to their specificity and self-replication, phages provide therapy advantages. Yet, any successful application of the system calls for accurate prediction of the interaction between a phage and its host. Culture-based methods are both lengthy and restricted in scope. Computational models based on genomic data have therefore become essential in the quest for efficiently predicting PHIs. This will increase efficiency by enabling the models to pre-select candidates that then do the experimentation **[1, 2, 3, 4, 5]**.

First, the sequence-based homology searches: Identifying conserved motifs and domains that mediate phage-host recognition.

Second, the machine learning algorithms: Predictive models can be trained on large datasets of known phage-host interactions (PHIs), using features derived from genomic sequences, phage infection patterns, and structural information. Crucially, integrating experimentally validated host range matrices — which reflect infectivity patterns across diverse phage and bacterial strains — can significantly enhance model performance. This combined approach enables more accurate predictions of novel phage-host interactions and supports the rational selection of phages for therapeutic applications. Network-based approaches: Constructing interaction networks based on genomic proximity and co-occurrence patterns, revealing potential phage-host associations.

These computational approaches are not intended to replace experimental validation entirely. Rather, they serve to prioritize and streamline experimental efforts, enabling researchers to focus on the most promising phage candidates. Combining information on phage infectivity patterns against specific pathogen collections with phage and pathogen genomic information will contribute to predicting phage infection outcomes on new pathogen isolates and thus support a more efficient selection of phages against changing pathogen communities. The advent of high-throughput sequencing has generated vast genomic datasets, demanding scalable computational approaches for analyzing phage-host interactions **[6]**. Machine learning (ML)-based tools have emerged as powerful solutions for predicting these interactions, offering the potential to rapidly identify suitable phage candidates for various applications.

However, a critical challenge remains: the predictive performance of ML tools varies substantially across taxonomic levels, from broad phyla to specific species **[7]**. This inconsistency complicates tool selection for targeted research questions and can hinder the reproducibility and reliability of computational predictions **[9,10]**. Despite the proliferation of available tools, a systematic evaluation of their effectiveness across the taxonomic hierarchy has been lacking, leaving researchers without clear guidelines for optimal tool choice **[11]**.

To address this gap, we conducted a systematic review and meta-analysis of ML-based phage-host interaction prediction tools. Our study evaluates tool performance across diverse taxonomic categories, employing robust statistical methods, including meta-regression and precision consistency tests **[12,13,14]**. By rigorously assessing predictive capabilities and identifying factors influencing variability **[15]**, our findings provide critical insights into the strengths and limitations of current tools. This work offers practical guidance for researchers, enabling informed selection of the most appropriate tool based on specific research needs and desired taxonomic resolution.

## Materials and methods

This systematic review and meta-analysis were conducted following the PRISMA (Preferred Reporting Items for Systematic Reviews and Meta-Analysis) guidelines **[16]**. The process involved five key stages: (1) planning and protocol development; (2) comprehensive literature search and retrieval; (3) study selection and eligibility assessment; (4) data extraction and quality assessment; and (5) data synthesis and meta-analysis.

### Search strategy and study selection

A systematic literature search was performed across three electronic databases: PubMed, Scopus, and Web of Science, covering the period from 2014 to 2025. The search strategy combined relevant keywords and controlled vocabulary terms related to phage-host interactions and machine learning-based prediction tools. The specific search terms included: (“phage” OR “bacteriophage”) AND (“bacteria”) AND (“phage host interaction”) AND (“machine learning”).

The search results were initially screened by title and abstract to identify potentially eligible studies. Full-text articles of these studies were then retrieved and assessed for eligibility based on pre-defined inclusion and exclusion criteria.

### Inclusion criteria

Studies were included if they:

1. Focused on the development or evaluation of machine learning-based tools for predicting phage-host interactions.
2. Used empirical data to validate the performance of the prediction tools.
3. Were published in peer-reviewed journals, conference proceedings, or book chapters.
4. Were published between 2014 and 2024.

### Exclusion criteria

Studies were excluded if they:

1. Did not involve machine learning-based prediction of phage-host interactions.
2. Focused on other types of biological interactions or prediction tasks.
3. Data extraction and quality assessment.

Data from eligible studies were independently extracted by three reviewers using a standardized data extraction form. Discrepancies between reviewers were resolved through discussion and consensus. The extracted data included information on:

1. Study characteristics (e.g., authors, year of publication, study design).
2. Prediction tools (e.g., name, algorithm, features).
3. Datasets used for training and testing.
4. Performance metrics (e.g., accuracy, Proportion of Correctly Predicted PHIs, precision).
5. Taxonomic levels of analysis.

The methodological quality of the included studies was assessed using the Newcastle-Ottawa Scale for non-randomized studies **[17]**.

### First keyword screening: search in titles, abstracts, and keywords sections

Scopus, WoS, and PubMed (Medline) databases were searched with the following search words: (phage) OR (bacteriophage) AND (bacteria) AND (Phage host interaction) in titles, abstracts, and keywords of the publications. Database searches included documents published since 1960 for Scopus, and from 2014 and 2024.

### Second keyword screening: search in the materials and methods section

Keywords were searched in the materials and methods section of the publications with the following words: (phage) OR (bacteriophage) AND (bacteria) AND (Phage host interaction).

#### Document review

A full-text manual review was conducted by three independent reviewers according to the inclusion and exclusion criteria.

#### Selection criteria

Eligibility criteria were based on the **PICO** approach (Population: Tools of machine learning employed for identification of phage host interaction, exposure: phage host interaction in taxonomic levels using the predictive tools, the comparison group: accuracy of the different predictive tools, and several taxonomic levels, and outcome: % of Proportion of Correctly Predicted PHIs of the taxonomic level).

**Figure 1**. Systematic review and meta-analysis: Machine learning programs used to identify phage host interaction

**Figure 1.**
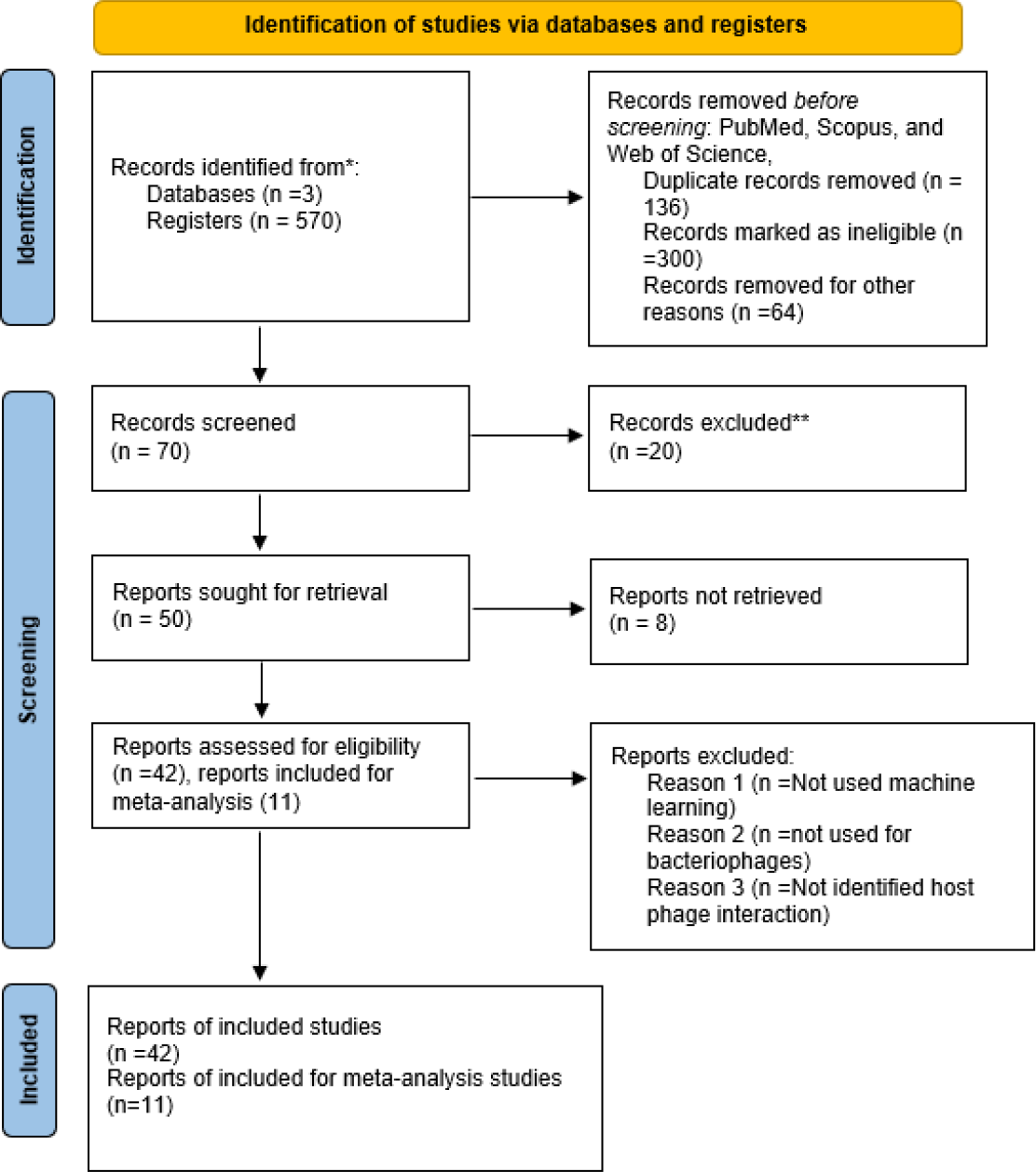
PRISMA flow diagram of the study selection process. Of 570 records identified across PubMed, Scopus, and Web of Science, 42 studies met inclusion criteria for the systematic review and 11 provided sufficient quantitative data for meta-analysis (61 datasets total).

### Data analysis

From each study, we extracted the proportion of experimentally validated phage-host interactions (PHIs) correctly predicted by the evaluated tools. Since not all studies reported true negatives or false positives, we focused on recall (i.e., the fraction of true interactions recovered), rather than overall accuracy.

A Meta-analysis was performed using a random-effects model to estimate the pooled performance of machine learning-based phage-host interaction prediction tools across different taxonomic levels. Subgroup analyses were conducted to explore potential sources of heterogeneity, such as the type of machine learning algorithm, the features used for prediction, and the taxonomic level of analysis using the R project. Besides, an ANOVA was performed to determine if there is a significant difference between taxonomical level identification and prediction tools.

## Results

### Systematic Review and Study Selection

The systematic search across PubMed, Scopus, and Web of Science databases, using the pre-defined search approach, identified 570 records. After removing duplicates and screening titles and abstracts, 70 potentially relevant articles were retrieved for full-text review. Following application of the eligibility criteria (detailed in the Methods section), the 42 studies included in the systematic review were selected based on predefined inclusion criteria, and 11 of them provided sufficient quantitative data (such as precision, size of genome of bacteria and phages values) to be included in the meta-analysis. A PRISMA flow diagram (Figure 1) visually summarizes the study selection process.

The values analyzed reflect results originally reported by the authors in each study. We did not reanalyze raw data, but rather extracted and compared performance metrics as reported in the published results.

Furthermore, the **Proportion of Correctly Predicted PHIs**, as defined by each study. We standardized these metrics to make them comparable across tools or taxonomic levels, and further details can be found in the Methods and Results sections.

### Meta-Analysis: Phage-host interactions prediction tools have been evaluated for performance across all taxonomic ranks

This first meta-analysis has been carried out to evaluate the performance of machine learning-based phage-host interaction (PHI) prediction tools or methods evaluated across Domains, Phyla, Classes, Orders, Families, Genera, and Species. The figures reported are recall values, representing the proportion of experimentally validated PHIs that have been correctly predicted by the tools under study. Therefore, there exist truly stark differences in predictive performance across these taxonomic levels analyzed, as shown in Figure 2.

**Figure 2.**
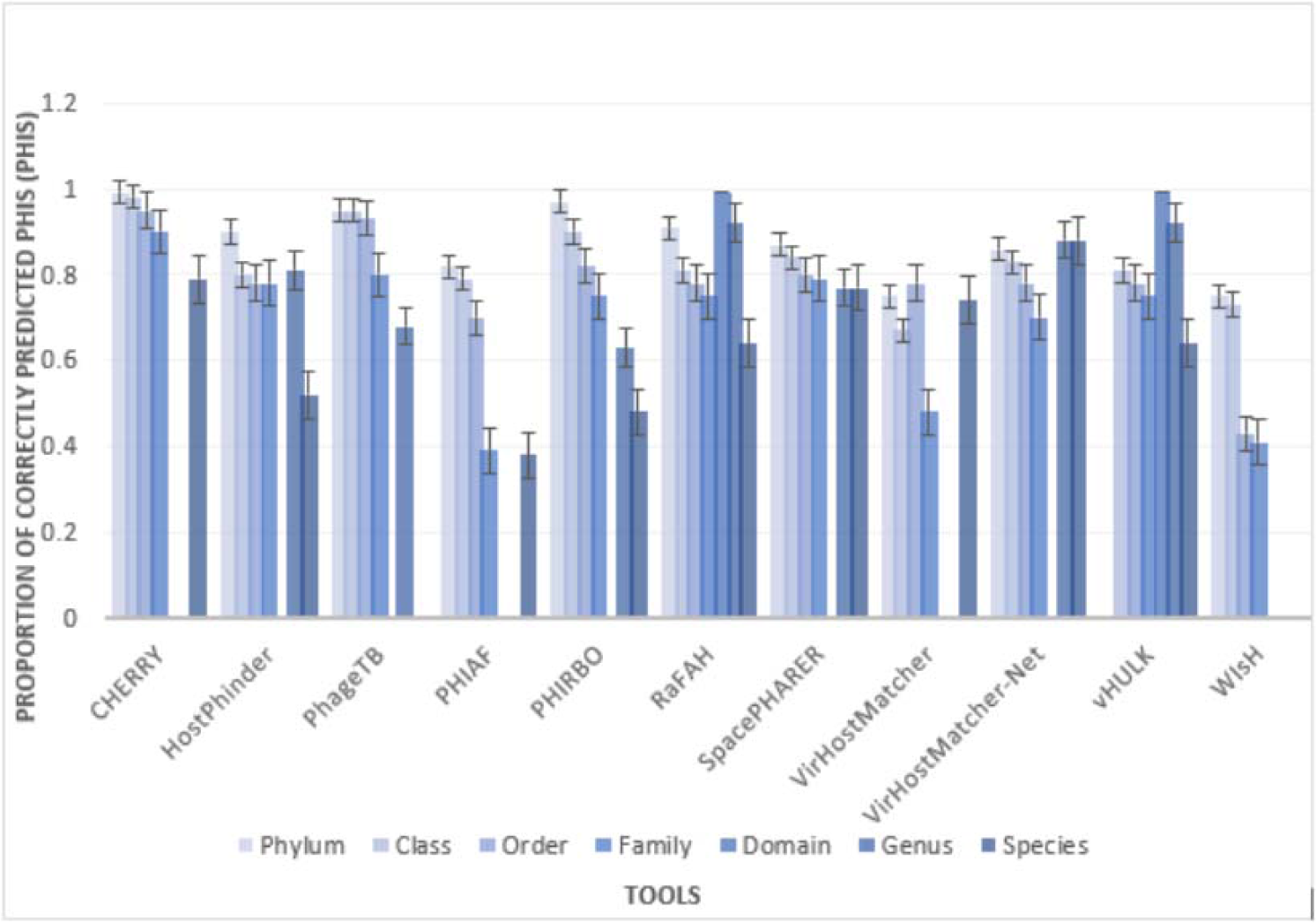
Recall (Proportion of Correctly Predicted PHIs) of machine learning tools across taxonomic levels (Domain, Phylum, Class, Order, Family, Genus, Species). CHERRY achieves the highest overall recall; VirHostMatcher-Net leads at Species level (0.88); WIsH consistently underperforms. Error bars represent variability across datasets.

Although CHERRY was consistently found to present favorable recall values (0.79 at Species; 0.90 at Family; 0.95 at Order; 0.98 at Class; 0.99 at Phylum) when considered across the taxonomic rank, choosing the best tool tended to be contingent upon the context. At the Genus rank, RaFAH and vHULK were able to compete with CHERRY, looking at the recall values (both 0.92). VirHostMatcher-Net, contrarily, took the lead in recall at the Species rank (0.88) over any other tools considered here. WIsH had the lowest recall value in almost all taxonomic ranks (e.g., Phylum 0.75, Class 0.73, Order 0.43, Family 0.41), with PHIAF having the lowest recall value only for Species.

**Figure 2. Proportion of Correctly Predicted PHIs** of different tools to detect the phage host interaction, in various levels as phylum, class, order, family, domain, genus, and species.

### Statistical Analysis of Precision Consistency Across Taxonomic Levels

To assess the consistency of precision across taxonomic levels, the Shapiro-Wilk test for normality and the ANOVA test (a non-parametric test) were performed. The results of the Shapiro-Wilk test indicated no substantial deviations from normality for any of the taxonomic levels. Furthermore, the ANOVA test revealed no statistically significant differences in precision distributions across the taxonomic groups (Table 1).

**Table 1:**
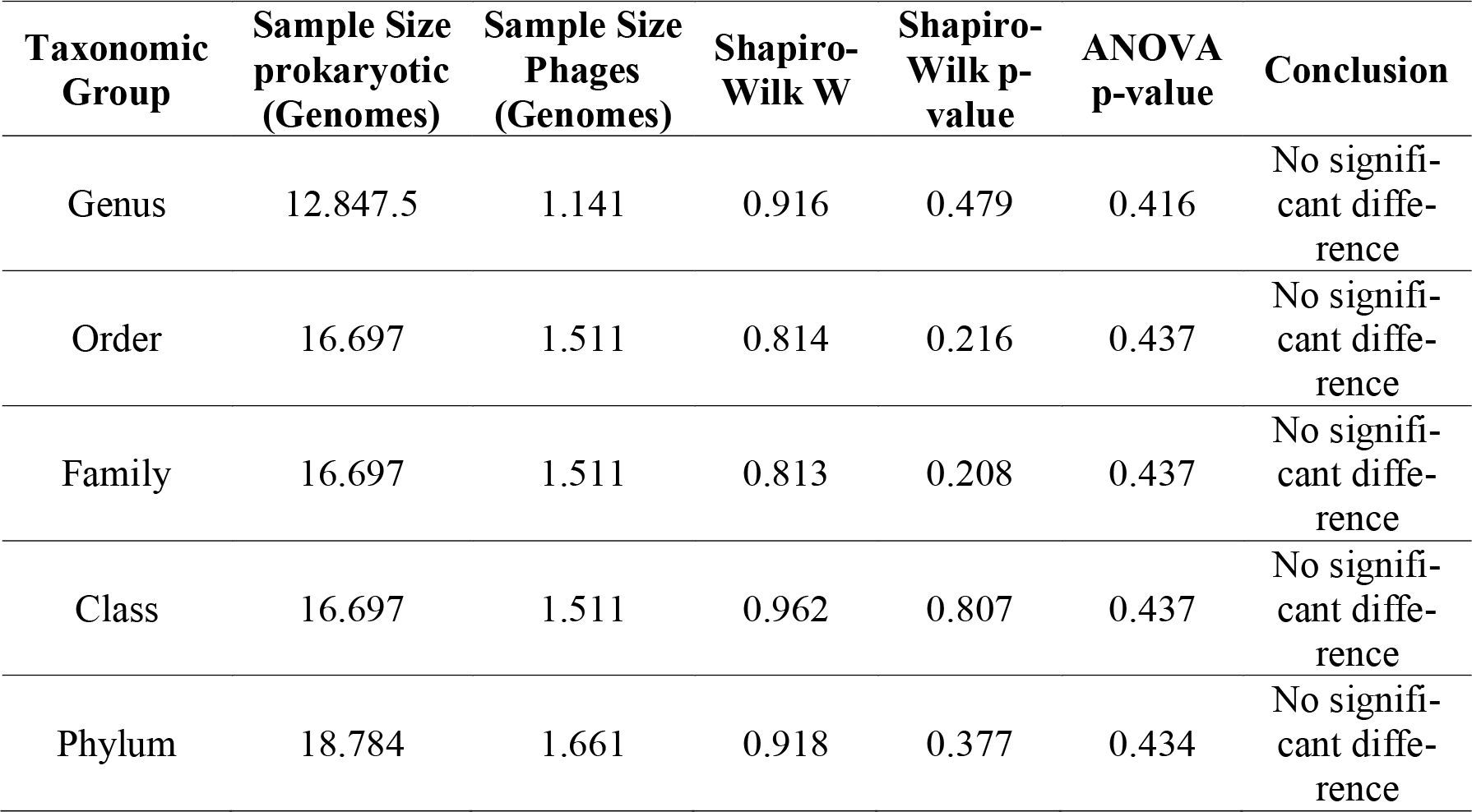
Comparison of phage-host interaction using different methods Across Taxonomic Levels.

These findings underscore the consistency of precision across taxonomic levels in both prokaryotic and phage genome predictions. This consistency highlights the robustness of the machine learning tools employed to identify phage-host interactions, demonstrating their reliability across multiple levels of taxonomic classification. The results support the suitability of these methodologies for broad applications in host-phage interaction studies.

### Meta-Regression Analysis of Subgroup Effects on Precision

A mixed-effects meta-regression model was used to investigate the influence of taxonomic sub-groups on prediction precision. The model included taxonomic level (Domain, Family, Genus, Order, Phylum, Species) as a categorical predictor variable. The analysis revealed statistically significant effects of specific taxonomic levels on prediction precision (Table 2). In the meta-regression model, each taxonomic level was included as an independent variable coded as a categorical effect.

**Table 2.** Meta-Regression Analysis of Subgroup Effects on Precision.

| Subgroup | Effect ($\beta$) | p-value | Significance | Interpretation |
| --- | --- | --- | --- | --- |
| Domain | 0.1195 | 0.1437 | Not significant | No evidence of a significant effect on precision. |
| Family | -0.1464 | 0.0006 | *** | Significant reduction in precision by 0.1464 for this subgroup. |
| Genus | -0.0653 | 0.1867 | Not significant | Does not significantly affect precision. |
| Order | -0.0527 | 0.2162 | Not significant | Similar to Genus, no significant impact. |
| Phylum | 0.0532 | 0.2258 | Not significant | Small increase but not significant in precision. |
| Species | -0.1944 | <0.0001 | *** | Significant reduction in precision by 0.1944. |

Meta-regression is used here to explore whether certain subgroup characteristics can systematically influence the outcome, in this case, precision. A random-effects model was employed to account for heterogeneity across studies. Table 2 presents the results of a meta-regression analysis conducted to examine the effect of different taxonomic levels (Domain, Phylum, Class, Order, Family, Genus, and Species) on the precision of host prediction.

Besides, each parameter (β coefficient) represents the estimated change in precision when the sub-group level is used as a reference, while controlling for variability among studies. A positive β indicates a potential increase in precision, whereas a negative β suggests a reduction. The p-values indicate the statistical significance of these effects.

Our findings show that among all levels tested, Family and Species levels have a statistically significant negative impact on precision (p < 0.001), with β values of −0.1464 and −0.1944, respectively. This suggests that using predictions at these taxonomic levels may lead to a greater drop in precision compared to other levels. Conversely, no significant effects were observed for Domain, Genus, Order, or Phylum, indicating that variations at these levels do not strongly influence prediction precision across studies.

The mixed-effects meta-regression model demonstrated a strong fit to the data, as indicated by a low Bayesian Information Criterion (BIC) of −13.3376. This low BIC value suggests a good balance between model complexity and explanatory power. The model’s statistical significance was confirmed (p < 0.0001), indicating a non-zero baseline level of Proportion of Correctly Predicted PHIs across the included studies. Critically, the model explained all observed heterogeneity in precision, as evidenced by a τ^2^ value of 0. This result strongly suggests that taxonomic level, the primary moderator included in the model, fully accounts for the variability in the predictive Proportion of Correctly Predicted PHIs observed across the studies. The I^2^ statistic of 0.00% further reinforces this conclusion, indicating that 100% of the heterogeneity was explained by the model.

These results highlight the critical role of methods and taxonomic subgroups as moderators in explaining the variability of the Proportion of Correctly Predicted PHIs in the studies. A bubble plot (Figure 3) illustrates the relationship between the effect size estimates and the moderators, providing a visual summary of the meta-regression results.

**Figure 3.**
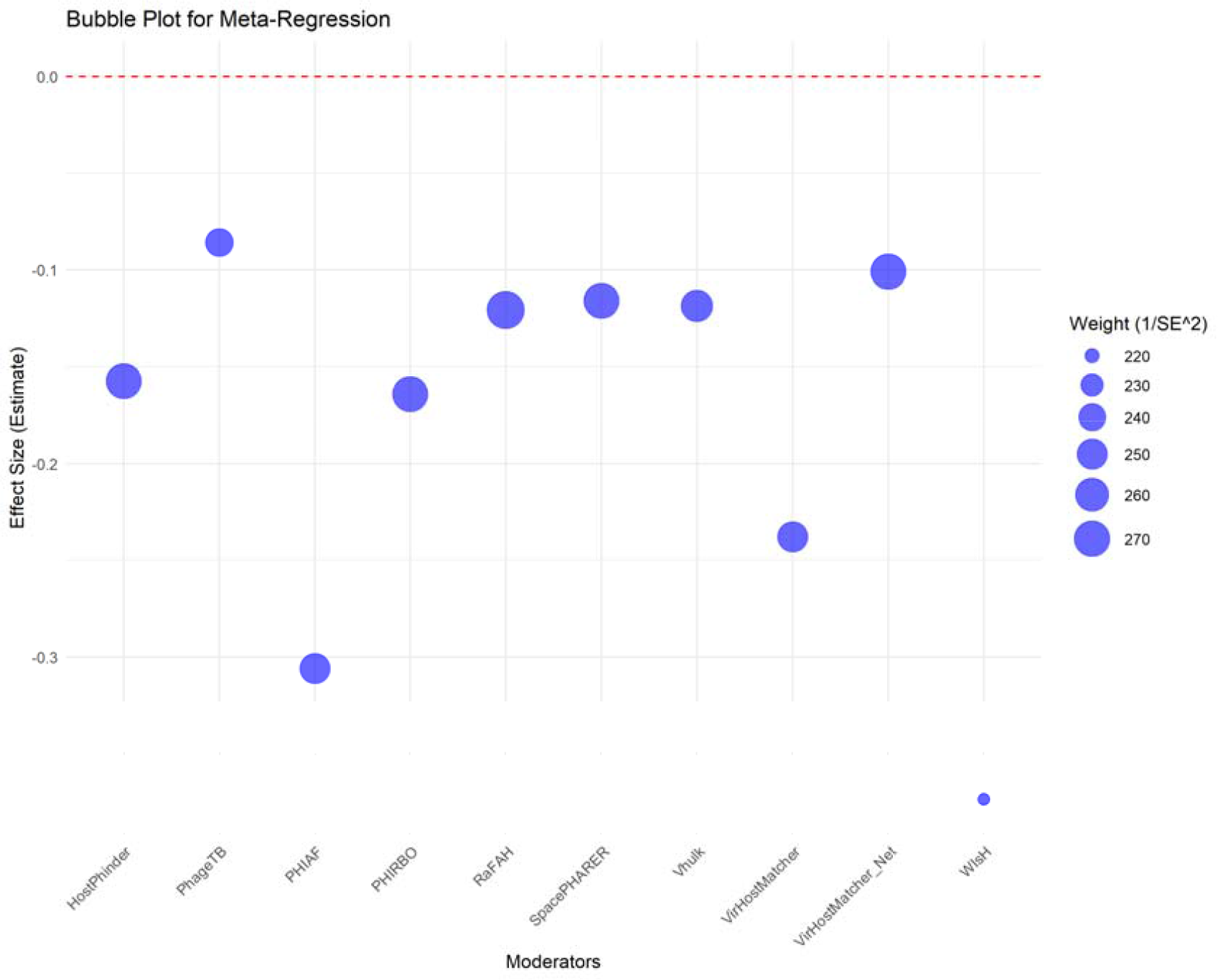
Bubble plot of meta-regression results showing the relationship between β-coefficient effect sizes and taxonomic moderators. Bubble size reflects study weight. Family (β = −0.1464, p = 0.0006) and Species (β = −0.1944, p < 0.0001) show statistically significant negative effects on prediction precision.

The findings emphasize the utility of mixed-effects models in uncovering nuanced relationships between the Proportion of Correctly Predicted PHIs and moderating factors, offering valuable insights into the performance and limitations of prediction methods across taxonomic levels.

**Figure 3**. Bubble plot of the relationship between the effect size estimates and the moderators.

## Discussion

This study highlights the potential and current limitations of using computational tools to support phage therapy. While phage therapy offers a highly specific alternative to antibiotics, its practical application depends on identifying effective phages for individual pathogens. Traditional methods depend on phenotypic susceptibility testing, which is time-consuming and dependent on pathogen isolation. In contrast, the bioinformatic tools evaluated in this study RaFAH, vHULK, and HostPhinder, offer a rapid, genome-based strategy to predict phage-host relationships. We found that predictions at higher taxonomic ranks, such as genus and family, are generally robust and consistent. Nevertheless, at the species and particularly strain level, predictive accuracy declines considerably, limiting their standalone utility in clinical settings where precision is essential. These results highlight the need to integrate computational predictions with targeted phenotypic assays in a hybrid workflow to accelerate phage selection while maintaining the accuracy required for therapeutic applications.

Numerous computational tools have been developed to predict phage-host interactions using genomic information. These tools rely on different underlying principles to infer host identity [18, 19]. Alignment-based tools such as HostPhinder [20] and SpacePHARER [21] use sequence similarity between the phage genome and known bacterial genomes, often based on shared k-mers or gene content. Others, like WIsH [22] and VirHostMatcher [23], utilize oligonucleotide frequency patterns (genomic signatures) to compare compositional similarity between phages and potential hosts. Tools such as CHERRY [24], RaFAH [25], vHULK [26], and PHIRBO [27] incorporate machine learning or deep learning models, trained on large datasets of known phage-host pairs, to detect complex genomic features associated with host specificity. PhageTB [28] use CRISPR spacer matches, BLAST alignments, and Hybrid machine learning, and PHIAF [29] combines multiple features, combines DNA sequence features and GAN-Based Data Augmentation. Figure 4 summarizes the main input data, computational methods, and predictive outputs employed by current phage–host prediction tools. Approaches include genomic similarity, k-mer-based composition analysis, and AI-driven models to infer host taxonomy and guide phage therapy applications (Figure 4).

**Figure 4.**
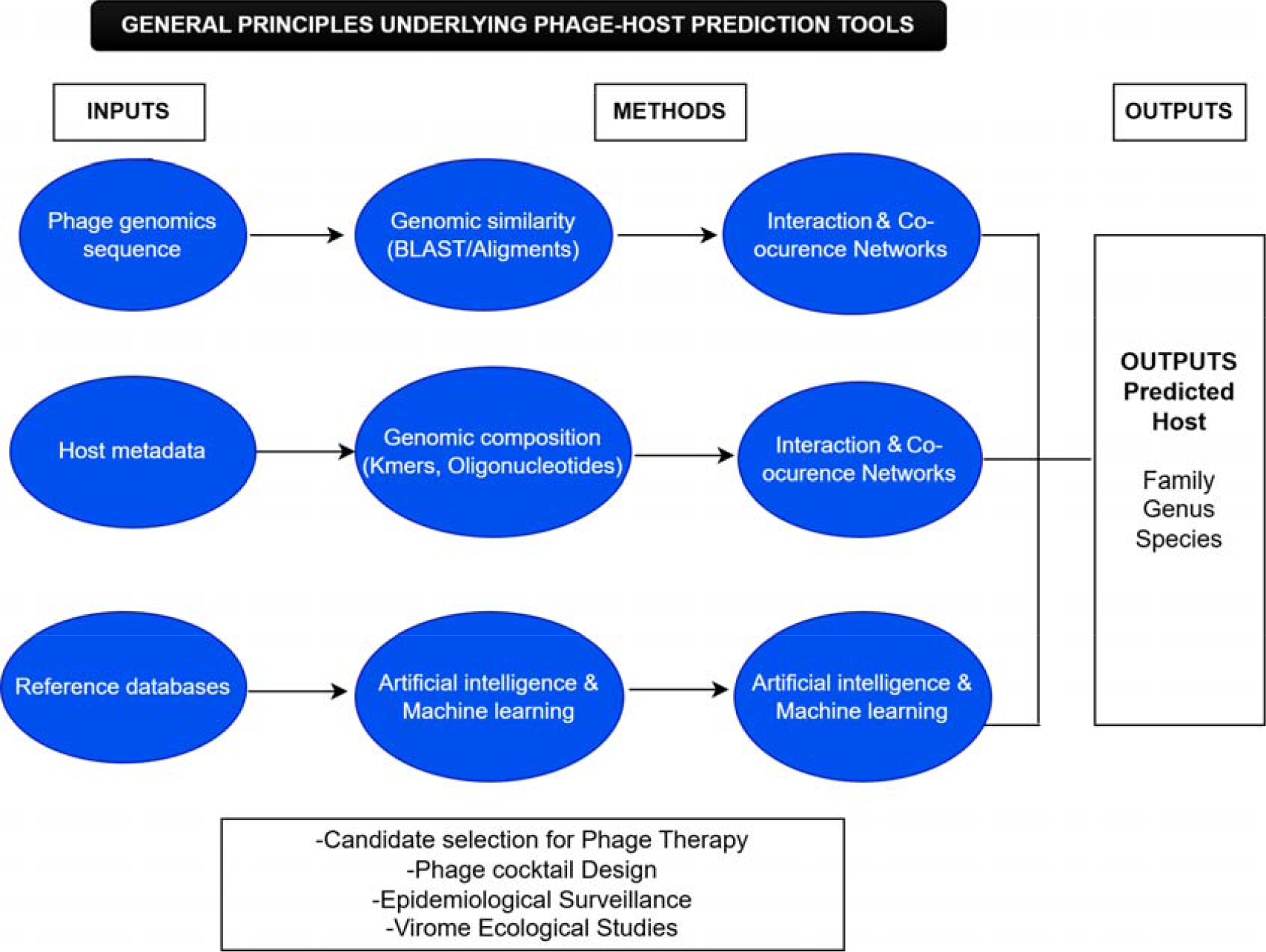
Overview of core computational approaches for phage–host interaction prediction. Tools are classified by input data type (genomic similarity, k-mer composition, ML/deep learning, CRISPR spacers, hybrid) and predictive scope (taxonomic resolution and application context).

**Figure 4**. Overview of Core Computational Approaches for Phage–Host Interaction Prediction Tools These tools differ not only in methodology but also in their predictive scope: while some are optimized for host predictions at higher taxonomic levels (e.g., family or genus), others aim to achieve finer resolution, ideally down to species or strain level. However, accuracy tends to decrease as taxonomic resolution increases, limiting their standalone use in clinical phage therapy where strain-specific identification is crucial.

In this study, we applied a subset of these tools, including WIsH, HostPhinder, and RaFAH to evaluate their performance on a curated dataset of phage genomes with known bacterial hosts, compiled from published scientific articles. These tools represent different methodological classes (compositional, alignment-based, and machine learning). We observed that while predictions at the genus and family levels were relatively consistent across tools, the accuracy significantly declined when attempting to resolve host identity at the species level. This limitation is critical in the context of phage therapy, where strain-specific targeting is essential. Currently, no tool reliably predicts host at the strain level without complementary phage-susceptibility validation. Computational predictions can serve as a first-pass screening step, narrowing down candidate phages for in vitro testing. Regarding molecular targets, the tools typically infer host specificity indirectly by detecting shared genomic features, such as CRISPR spacers, tRNA genes, or nucleotide composition signatures that reflect past evolutionary interactions. Though these are substitutions rather than direct molecular interactions (e.g., phage tail fiber–receptor binding), which remain challenging to predict solely from sequence data. Hence, while these tools are valuable for guiding host-range hypotheses, their integration into therapeutic pipelines requires careful interpretation and experimental validation.

### Variability in the performance of prediction tools

The meta-analysis further elucidates inherent paradoxes in tool performances, as the latter are highly dependent on the methodological underpinnings of each approach. As for CHERRY, this tool’s capability, especially across broader taxonomic levels, is dictated by the inherently versatile deep learning architecture that it possesses. Likewise, RaFAH and vHULK, basing their analyses on random forest algorithms coupled with alignment-free features, can maintain competitive performances at the genus level by grasping larger-scale evolutionary depths. However, VirHostMatcher-Net, which captures oligonucleotide frequencies and conducts network-based analyses, attains better performance at a species-level resolution, where genetic subtleties are critical. WIsH attempted to mark the model à la k-mer, yet it could never work well across the taxonomic scales because of its lack of deep consideration of the complex biological processes. For example, PHIAF, an initially promising alignment-free method, may have suffered from feature overfitting at the species level in its particular implementation. These subtle differences in methodology and their resultant advantages and disadvantages are relevant to rational tool selection and are briefly tabulated in Supplementary Table S1 for all tools considered. Deliberate consideration of these methodological differences not only informs appropriate tool selection but also indicates where enhancements may be targeted to enhance the accuracy and reliability of prediction over the full range of taxonomic scales.

### Impact of taxonomic resolution on prediction: Proportion of Correctly Predicted PHIs

The analysis of the Proportion of Correctly Predicted PHIs in taxonomic hierarchies revealed distinct patterns. The broader taxonomic categories (domain, phylum, and species) showed a high Proportion of Correctly Predicted PHIs and consistency. Domain-level predictions, in particular, achieved a near-perfect Proportion of Correctly Predicted PHIs (0.99). However, a clear trend of decreasing Proportion of Correctly Predicted PHIs emerged as the taxonomic resolution became finer. The average Proportion of Correctly Predicted PHIs at the family and order levels (0.682 and 0.775, respectively) was significantly lower than that observed at the broader levels. This decrease in performance at finer taxonomic resolutions aligns with findings from previous studies **[30]**. These studies have highlighted the increased complexity of distinguishing between closely related taxa at finer levels, emphasizing the need for improved databases and algorithmic approaches. Our findings further corroborate this trend, demonstrating that while broader classifications are often reliable, finer-level predictions remain a significant challenge. The greater variability at the family and order levels likely reflects the greater diversity and complexity of phage-host interactions at these finer resolutions.

Several studies have found that forecasts at higher taxonomic levels are frequently more reliable, although finer-level predictions (e.g., at the species or strain level) remain a significant difficulty. This greater variability at finer resolutions is most likely owing to the inherent complexity and diversity of phage-host interactions, and partly driven by their co-evolution. While genetic techniques have produced useful insights, they alone are insufficient to capture the full range of these interactions.

To solve this shortcoming, we propose combining prediction methods with phenotypic data to increase their accuracy. Linking phenotypic traits of phage host range and host susceptibility patterns to the genetic variation and determinants of phage-host interactions should considerably improve our ability to make precise predictions at fine taxonomic resolutions. Phenotypic tests, including highthroughput infectivity screens and host resistance profiling, should therefore be combined with genomic data to discover critical factors influencing phage-host interactions.

Furthermore, we propose the creation of machine learning models that use both genomic and phenotypic data. Such integrated models would allow for the discovery of intricate and hidden links between genetic variation and phenotypic features, which is critical for boosting prediction accuracy, especially at lower taxonomic levels.

Finally, this integrative method should be accompanied by ongoing experimental validation to guarantee that computational predictions are correct and dependable. An iterative procedure that combines in silico predictions with empirical data will improve the predictive capacity of these tools while also deepening our understanding of phage-host dynamics. Incorporating both genomic and phenotypic data will be critical for overcoming present constraints and making therapeutically relevant predictions in phage therapy and related applications **[31]**.

### Statistical consistency and meta-regression analysis

While the Shapiro-Wilk and ANOVA tests did not reveal significant differences in the Proportion of Correctly Predicted PHIs distributions between taxonomic groups, suggesting a degree of overall consistency in tool performance, the meta-regression analysis provided more nuanced information. This analysis confirmed the significant moderating effect of the taxonomic subgroup on the prediction Proportion of Correctly Predicted PHIs. Specifically, the family and species levels were associated with significant reductions in the Proportion of Correctly Predicted PHIs. These findings underscore the particular challenges associated with predicting interactions at these finer taxonomic resolutions. The meta-regression model explained all the heterogeneity observed in the Proportion of Correctly Predicted PHIs (τ^2^ = 0, I^2^ = 0.00%), demonstrating the strong influence of the taxonomic level on the prediction of the Proportion of Correctly Predicted PHIs. This suggests that the taxonomic level is the main driver of variability in prediction Proportion of Correctly Predicted PHIs between studies.

### Implications for future research and tool development

Our findings have significant implications for future research and tool development in the field of phage-host interaction prediction. First, the variability observed in tool performance at the taxonomic levels calls for continued efforts to refine prediction algorithms, particularly for finer taxonomic categories. Future research should focus on the development of specialized tools and databases tailored to the specific challenges of predicting interactions at the family and order levels. Second, the significant moderating effect of the taxonomic subgroup highlights the need for a deeper understanding of the biological factors that influence phage-host specificity at different taxonomic resolutions. The integration of taxon-specific characteristics in prediction models could significantly improve their Proportion of Correctly Predicted PHIs. For example, recent studies have explored the use of protein embeddings **[32]** and graph contrastive augmentation **[33]** to improve phage-host interaction prediction, demonstrating the potential of incorporating advanced computational techniques.

Researchers should carefully consider the desired taxonomic resolution when selecting a prediction tool. While established methods may be sufficient for broader classifications, finer-level predictions require more specialized approaches. The prediction of phages for phage therapy applications has to identify phage-host pairs that match at the strain level. Our results, therefore, emphasize the need for continuous development and validation of prediction tools, particularly at a high-resolution taxonomy, to ensure reliable and accurate predictions of phage-host interactions. This could involve the exploration of new machine learning architectures **[34]**, the incorporation of multidimensional biological information, or the use of transfer learning techniques **[35]**.

In phage therapy applications, prediction resolution must often extend beyond the species level to capture intra-species variation. Such strain-level differences are critical for ensuring therapeutic efficacy, as phage infectivity can vary substantially among strains within the same species. This represents a fundamental limitation of current computational tools, which frequently fail to account for intra-species diversity. Addressing this gap requires that future prediction frameworks be explicitly designed to resolve strain-level variation. We propose the following approaches to improve prediction precision at these finer resolutions:

1. Integration of Strain-Level Data: Prediction tools must incorporate strain-level genomic and phenotypic information to account for within-species heterogeneity. This requires high-resolution sequencing combined with detailed phenotypic characterization to improve prediction accuracy at the individual strain level. There is a critical need for methods capable of accurately predicting strain-level phage-host interactions across diverse bacterial genera, particularly under data-limited conditions **[36]**.
2. Advances in Machine Learning Architectures: The refinement of machine learning models, particularly through deep learning approaches, may enable the identification of subtle patterns underlying intra-species variation. Such models should be trained on large datasets that include strain-specific information to improve predictive resolution at the strain level.
3. Transfer Learning: Applying transfer learning approaches, where knowledge derived from one species or strain is extended to others, may improve the generalizability and scalability of predictive models, particularly for taxa with limited training data.
4. Integration of phage-host infection matrices with genetic determinants of infection:

Phenotypic assays that evaluate infectivity and resistance at the strain level are essential for advancing phage host range prediction. These assays should complement genomic data to provide a more comprehensive understanding of the factors governing phage-host interactions. Nevertheless, many functional and ecological complexities remain unaccounted for by current tools, even as their accuracy and taxonomic coverage continue to improve.

According to experimental evidence, such as that in Middelboe et al. **[37]**, phage pressure on marine bacterial populations quickens the rates of strain-level diversification in hosts’ range and susceptibility. This evolutionary dynamism is an immediate challenge for in silico models, which often assert static associations solely based on genomic similarity.

Besides, it has been shown **[38]** that prophage-encoded auxiliary metabolic genes, such as chitinases, can significantly alter host physiology and viability, forcing these organisms to survive under extreme conditions. This insinuates that functional interactions beyond mere molecular recognition must be contemplated in the following generation of predictive models. In turn, the same findings suggest that ecological and functional knowledge must be incorporated to improve and design sturdy prediction tools.

## Conclusion

This systematic review and meta-analysis provide a comprehensive assessment of the performance and variability of phage-host interaction prediction tools across taxonomic levels. Our findings underscore the importance of considering taxonomic resolution when selecting and developing prediction tools. By highlighting the strengths and limitations of current methodologies, this study contributes to a more informed and targeted approach to predicting phage-host interactions, ultimately advancing our understanding of phage biology and facilitating the development of effective phage-based therapies and biotechnological applications.

## Acknowledgments

MM was supported by grants from the European Union under the Horizon Europe Programme, Grant Agreement No. 101084204 (Cure4Aqua) and from the Innovation Fund Denmark project No. 2105-00014B (AQUAPHAGE). The authors thank Professor Robert Schooley for his insightful contribution to the discussion.

## Data Availability Statement

The data supporting the findings of this study are available within the article and its supplementary information files.

